# Broad-Spectrum, Patient-Adaptable Inhaled Niclosamide-Lysozyme Particles are Efficacious Against Coronaviruses in Lethal Murine Infection Models

**DOI:** 10.1101/2020.09.24.310490

**Authors:** Ashlee D. Brunaugh, Hyojong Seo, Zachary Warnken, Li Ding, Sang Heui Seo, Hugh D.C. Smyth

## Abstract

Niclosamide (NIC) has demonstrated promising *in vitro* antiviral efficacy against SARS-CoV-2, the causative agent of the COVID-19 pandemic. Though NIC is already FDA-approved, the oral formulation produces systemic drug levels that are too low to inhibit SARS-CoV-2. As an alternative, direct delivery of NIC to the respiratory tract as an aerosol could target the primary site of for SARS-CoV-2 acquisition and spread. We have developed a niclosamide powder suitable for delivery via dry powder inhaler, nebulizer, and nasal spray through the incorporation of human lysozyme (hLYS) as a carrier molecule. This novel formulation exhibits potent *in vitro* and *in vivo* activity against MERS-CoV and SARS-CoV-2 and may protect against methicillin-resistance staphylococcus aureus pneumonia and inflammatory lung damage occurring secondary to CoV infections. The suitability of the formulation for all stages of the disease and low-cost development approach will ensure wide-spread utilization

---

The Coronavirus Disease 2019 (COVID-19), caused by the Severe Acute Respiratory Syndrome-associated coronavirus 2 (SARS-CoV-2), was declared a pandemic by the World Health Organization (WHO) in March 2020(1). A rapid and continuing rise in cases worldwide followed this declaration which has overwhelmed healthcare systems(2, 3). At present, there are no widely available vaccines or treatments for coronaviruses (4, 5), and outbreaks are expected to continue potentially into 2025 (6). To address the ongoing pandemic, it is necessary that treatments are made rapidly available and at low-cost to patients and clinicians worldwide. One mechanism for achieving this has been to assess existing drugs for their utility against SARS-CoV-2. Recently, a screening of 48 FDA-approved drugs revealed that a compound called niclosamide (NIC), which has been historically utilized to treat intestinal tapeworms, also exhibits highly potent activity against SARS-CoV-2 in a Vero cell infection model(7). NIC has been previously demonstrated to enhance autophagy in Middle East respiratory syndrome coronavirus (MERS-CoV) infections through inhibition of SKP2(8), and it is hypothesized that a similar mechanism of action may enable its anti-SARS-CoV-2 activity(7). The broad-spectrum pharmaceutical utility of NIC appears to be related to its protonophoric activity, which results in disruptions of membrane pH gradients that may cause alterations in several signaling pathways. (9–18) The drug’s non-specific mechanism of action is illustrated by its efficacy against both gram-positive and gram-negative bacteria, including *Staphylococcus aureus* (19–21), *Pseudomonas aeruginosa* (22–24), *Acinetobacter baumannii* (22, 25), *Mycobacterium tuberculosis* (26, 27), and various bacterial biofilms (20, 21, 28), which may enable protection against secondary bacterial pneumonias associated with COVID-19. Furthermore, NIC has been noted to inhibit inflammatory cytokine release in lung tissues *in vivo*(29), which may provide an important protective effect against cytokine storm and acute respiratory distress syndrome (ARDS), one of the most feared sequela of COVID-19.

In terms of pharmaceutical development, a major limitation of NIC as an antiviral therapy stems from its poor solubility in water, which is reported at 1.6 mg/L(30). This makes systemic absorption of the drug by the oral route of administration at therapeutically relevant concentrations difficult. Administration of the existing FDA-approved chewable tablet formulation was found to result in inadequate systemic concentrations for inhibiting SARS-CoV-2 replication.(31, 32) As an alternative, direct delivery of niclosamide to the lung could overcome the limitations of the oral NIC formulation by generating high drug concentrations at the site of infection. For use against *P. aeruginosa*, Costabile *et al* previously developed an inhalable dry powder consisting of NIC nanocrystals embedded in mannitol particles.(33) However, these particles required a high amount of polysorbate 80 to ensure production of a stable suspension (10% w/w to NIC) which is beyond what is in currently FDA approved inhaled products (34). Furthermore, the utilization of mannitol as the carrier system may induce bronchospasm and cough(35–37) which may contribute to increased risk of spread of SARS-CoV-2 through generation of aerosolized respiratory droplets from infected individuals.

An additional considerable challenge in developing therapies for COVID-19 is the variable presentation of illness. Patients may act as asymptomatic carriers of the virus or develop severe pneumonias and acute respiratory disease(38), which can result in the requirement for mechanical ventilation. In the case of ventilated patients, a nebulizer is often used to delivery aerosolized drug to the lungs. However, for treatment of asymptomatic carriers or in regions of developing economies with reduced access to clean water sources, nebulizer-based aqueous products may present an undue burden and reduce therapy compliance. For these populations, dry powder inhaler (DPI) or nasal spray systems, or a combination of both, would be the preferred option based upon the rapid administration time and ease of use and could also potentially be utilized as a prophylactic therapy in high risk populations such as healthcare workers and first responders.

Utilizing repurposed NIC, and with the goal of developing a therapeutically effective, rapidly scalable and globally distributable antiviral therapy to reduce the spread of SARS-CoV-2, we describe an inhalable NIC formulation that can be administered using three major models or respiratory tract delivery systems: DPI, nasal spray and nebulizer. To achieve these aims, we utilized human lysozyme (hLYS), an endogenous protein in the upper and lower respiratory tracts, as a therapeutically active matrix material for the delivery of NIC to the airways, based upon its known anti-inflammatory (39), antiviral (40–43), and anti-bacterial activity (44–46) as well as its surface active properties. The antiviral, antibacterial, and anti-inflammatory efficacy of the NIC-hLYS powders were evaluated *in vitro* and *in vivo* in MERS-CoV and SARS-CoV-2 infection mouse models. The composition of the NIC-hLYS formulation was optimized using a constrained mixtures Design of Experiments (DoE) achieve particles with a size appropriate for inhalation both in the dry powder state and when reconstituted in aqueous media.

Physicochemical characterization of the optimized powder was performed and nasal, DPI, and nebulizer systems were developed and tested.

## RESULTS

### The addition of human lysozyme enhances *in vitro* niclosamide potency against coronaviruses

To determine the utility of hLYS as a carrier molecule for the nasal and pulmonary delivery of NIC, we assessed the *in vitro* antiviral activity of our novel formulation against a lysozyme-free NIC suspension (NIC-M). NIC-hLYS particles (0.7% w/w NIC) were administered at varying doses (based upon NIC content) to Vero E6 cells infected with MERS-CoV or SARS-CoV-2, and the EC50 was calculated based upon observed cytopathic effect (CPE). The addition of hLYS to the NIC formulation resulted in improved antiviral activity based upon reductions in the EC_50_ dose for MERS-CoV (0.016 μg/mL NIC to 0.0625 μg/mL NIC) and SARS-CoV-2 (0.030 μg/mL to 0.008 μg/mL) (Fig 1A and 1B).

**Figure 1:**
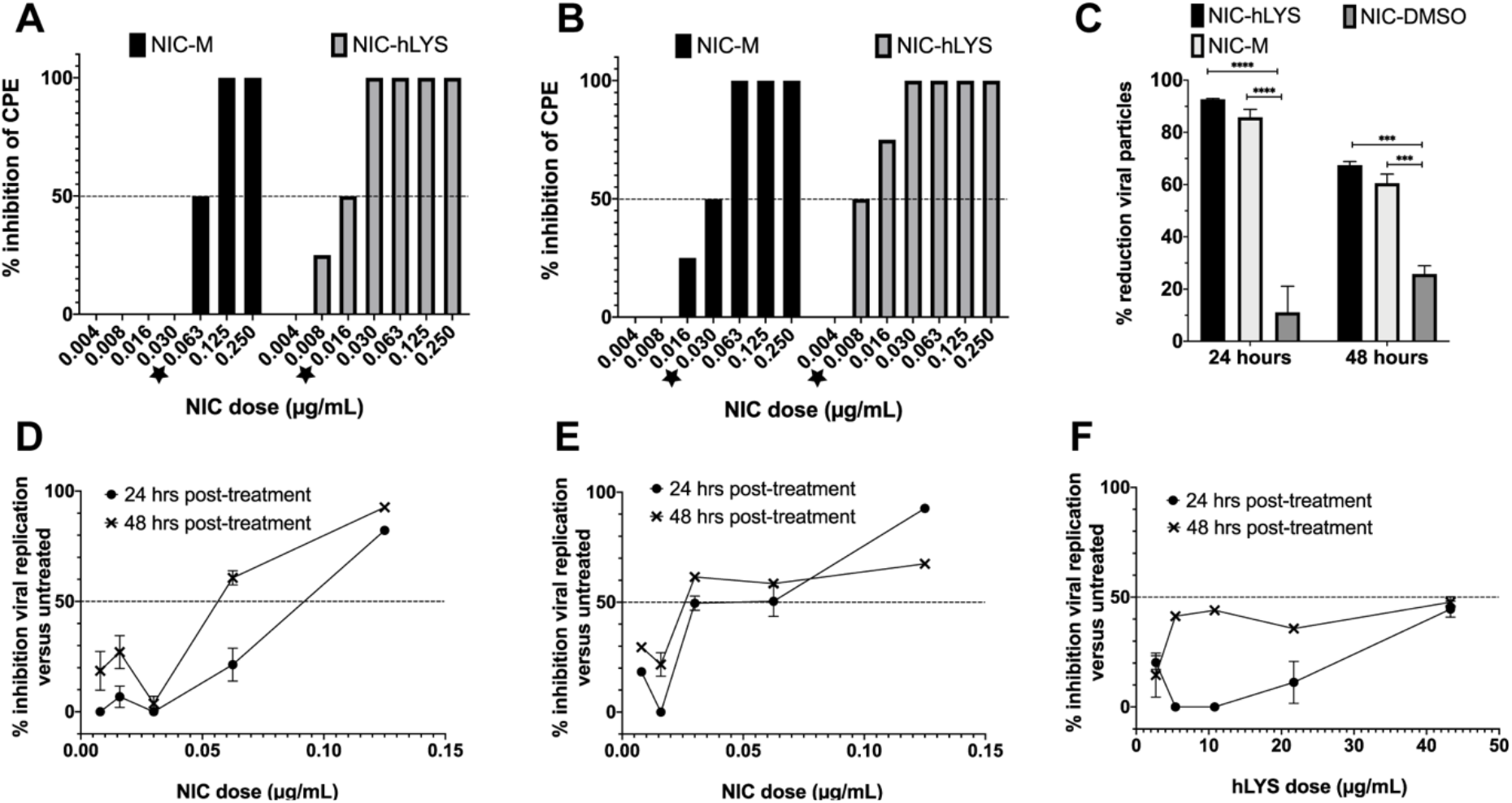
NIC-hLYS exhibited enhanced antiviral potency compared to NIC particles alone (NIC-M) in both MERS-CoV (A) and SARS-CoV-2 (B) infected cells. NIC-hLYS particles exhibited significantly higher inhibition of viral replication in SARS-CoV-2 infected Vero cells compared to an equivalent dose of solubilized NIC (NIC-DMSO), which indicates that improvements in solubility alone may not be sufficient to achieve maximal antiviral efficacy (C). Similar dose response profiles were noted for NIC-hLYS in both MERS-CoV (D) and SARS-CoV-2 (E), where an initial drop in activity is preceded by a sharp rise in activity. This profile was also noted in inhibitory assays for S. aureus. hLYS alone exhibits some antiviral activity which was more pronounced at 48 hours post dosing than 24 hours (F). Data presented as mean + SEM (n = 3). ***p < 0.001, ****p < 0.0001 using two-way ANOVA with Tukey’s multiple comparisons test.

We assessed the antiviral dose-response of NIC-hLYS particles in separate assay utilizing qPCR quantification of viral RNA collected from infected cells. At the highest dose tested (0.125 μg/mL NIC), Vero cells with an established MERS-CoV infection exhibited an 82.2% ± 0.8% decrease in viral load compared to untreated controls after 24-hours of exposure to NIC-hLYS particles (Fig 1D). This effect was sustained at 48 hours post-drug exposure (92.7% ± 2.0%; mean ± SD). A similar dose-response profile was achieved in Vero cells with an established SARS-CoV-2 infection (Fig 1E). A dose 0.125 μg/mL NIC resulted in a 24-hour inhibition of 92.7% ± 0.5% relative to untreated controls. However, in contrast to the MERS-CoV response, inhibition dropped to 67.5% ± 2.3% at 48 hours post-exposure in the SARS-CoV-2 infected cells. A separately conducted trypan blue viability assay in uninfected Vero E6 cells determined that the highest dose of NIC-hLYS utilized (0.125 μg/mL) had no effect on cell viability versus untreated controls (98.3% viability in treated cells).

Interestingly, hLYS alone appeared to exhibit activity against SARS-CoV-2 (Fig 1F), which has not been previously reported. Though the inhibitory activity was not as potent as that of NIC, it may contribute to observed increase in potency of NIC-hLYS in the CPE-based EC50 assay. NIC-hLYS, NIC-M and NIC dissolved in DMSO (NIC-DMSO) were compared for their inhibitory activity against SARS-CoV-2 at a NIC dose of 0.125 μg/mL. A two-way ANOVA with Tukey multiple comparisons test revealed statistically significant increase in viral inhibition of the NIC-hLYS particles versus NIC-DMSO at 24 hours (p < 0.0001) and 48 hours (p = 0.0001), though no significant differences were noted in the viral inhibition of NIC-hLYS and NIC-M formulations at the tested dose (Fig 1C), Thus, an improvement in solubility does not appear to be the mechanism for the increased activity of NIC-hLYS.

### Intranasal administration of NIC-hLYS particles to CoV-infected mice improves survivability and reduces viral loads in lungs, brain and kidneys

The *in vivo* efficacy of NIC-hLYS particles was assessed in lethal infection models for both MERS-CoV and SARS-CoV-2. HDDP4 transgenic mice were inoculated intranasally with 1 × 10^5^ pfu MERS-CoV and rested for 24 hours, after which once daily treatment with varying doses of intranasal NIC-hLYS (dosed based NIC component) was initiated (Fig 2A). In this initial efficacy study, animals were sacrificed at Day 6 to determine viral titres in brain and lung tissue compared to untreated controls. Notable decreases in brain viral titres was observed at the 120 μg/kg dose although these differences were not statistically significant (Two-way ANOVA with Dunnet’s multiple comparisons test, p = 0.1724) (Fig 2C). In a follow-up survival study utilizing lethal inoculum (1 × 10^5^ pfu), MERS-CoV-infected hDDP4 mice were dosed at 240 μg/kg NIC-hLYS daily by the intranasal route. By Day 10 (study endpoint), 43% of treated mice had survived compared to 0% of untreated controls (Fig 2B). Surviving mice were left untreated for an additional 3 days, at which point they were sacrificed. During this period of no treatment, the survival percentage remained at 43%. A statistically significant decrease in lung viral titres was noted in these surviving mice compared to the day 6 untreated controls from the earlier efficacy study (Two-way ANOVA with Dunnett’s multiple comparisons test, p = 0.011,). While brain viral titres did not exhibit further reduction from levels noted in the preliminary efficacy study, the inoculation of Vero E6 cells with viral particles obtained from lung and brain homogenates of surviving animals resulted in no observation of CPE at any of the inoculum concentrations tested, which indicates that remaining viral particles were not active. Thus, in the 43% of surviving animals it appears that the lethal MERS-CoV infection was essentially cured. Serological assays revealed that surviving animals expressed anti-MERS CoV IgG antibodies. Though notable in a lethal infection model, survival probability differences in treated and untreated mice were not statistically significant (Mantel-Cox test, p = 0.2984).

**Figure 2:**
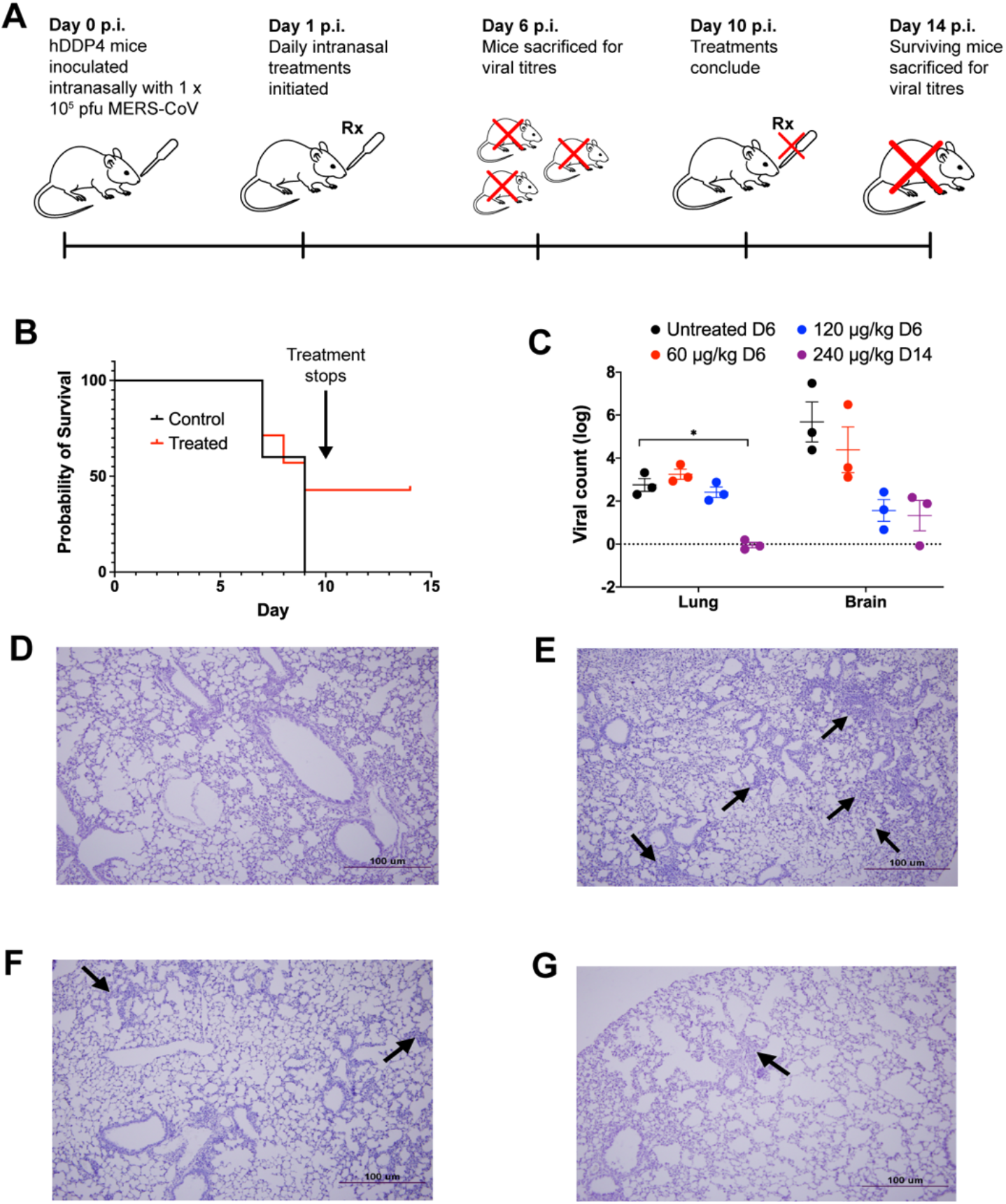
The efficacy of NIC-hLYS particles was assessed in a lethal MERS-CoV infection model (A). Once daily intranasal administration of NIC-hLYS particles suspended in 0.45% NaCl resulted in 43% survival in a lethal MERS-CoV infection (B) and produced a statistically significant decrease in lung viral titers at the highest dose tested (C). The viral particles obtained from lung and brain homogenates of surviving animals did not produce CPE when administered to Vero E6 cells, indicated that they were no longer active. Compared to lung tissue of uninfected mice (D), hDPP4-TG mice infected with MERS-CoV exhibited severe interstital pneumonia on Day 6 of infection (E), mice treated with NIC-hLYS exhibited milder iterstitial pneumona by Day 6 of treatment (F), which was further reduced by Day 14 (G). Data are presented as mean + SEM (n = 3). *p < 0.05, using two-way ANOVA with Dunnet’s multiple comparisons test.

In a similar study, the efficacy of NIC-hLYS particles was assessed in hACE2 transgenic mice infected intranasally with a lethal dose of SARS-CoV-2 (1 × 10^4^ pfu) (Fig. 3A). After a 24-hour rest period, daily intranasal treatment with 0.9% sodium chloride (untreated control) or NIC-hLYS (240 μg/kg NIC) was initiated. A portion of mice from each group (n = 3) were sacrificed at day 6 p.i. to determine viral loads in brain, kidney and lung tissues, while remaining animals were utilized in a 10-day survival study. Non-significant changes in day 6 viral titers were observed (Two-way ANOVA with Dunnet’s multiple comparisons test) (Fig. 3C). By the day 10 study endpoint, 30% of treated mice had survived, compared to 0% in the untreated arm (Fig 3B). Similar to the MERS-CoV study, these surviving mice were left untreated for an additional 3 days, at which point they were sacrificed (day 14 p.i.). The surviving mice exhibited a statistically significant decrease in viral loads in lung tissue compared to day 6 p.i. untreated controls (Two-way ANOVA with Dunnett’s multiple comparisons test), p = 0.023,), and no virus particles were detected in brain and kidney tissue using qPCR (Fig. 3C). Inoculation of Vero E6 cells with 10-fold dilutions of tissue homogenates resulted in no observed CPE, and surviving animals were sera positive for anti-SARS-CoV-2 antibodies. Similar to the MERS-CoV model, differences in survival probability between treated and untreated were not significant (Mantel-Cox test, p = 0.2959).

**Figure 3:**
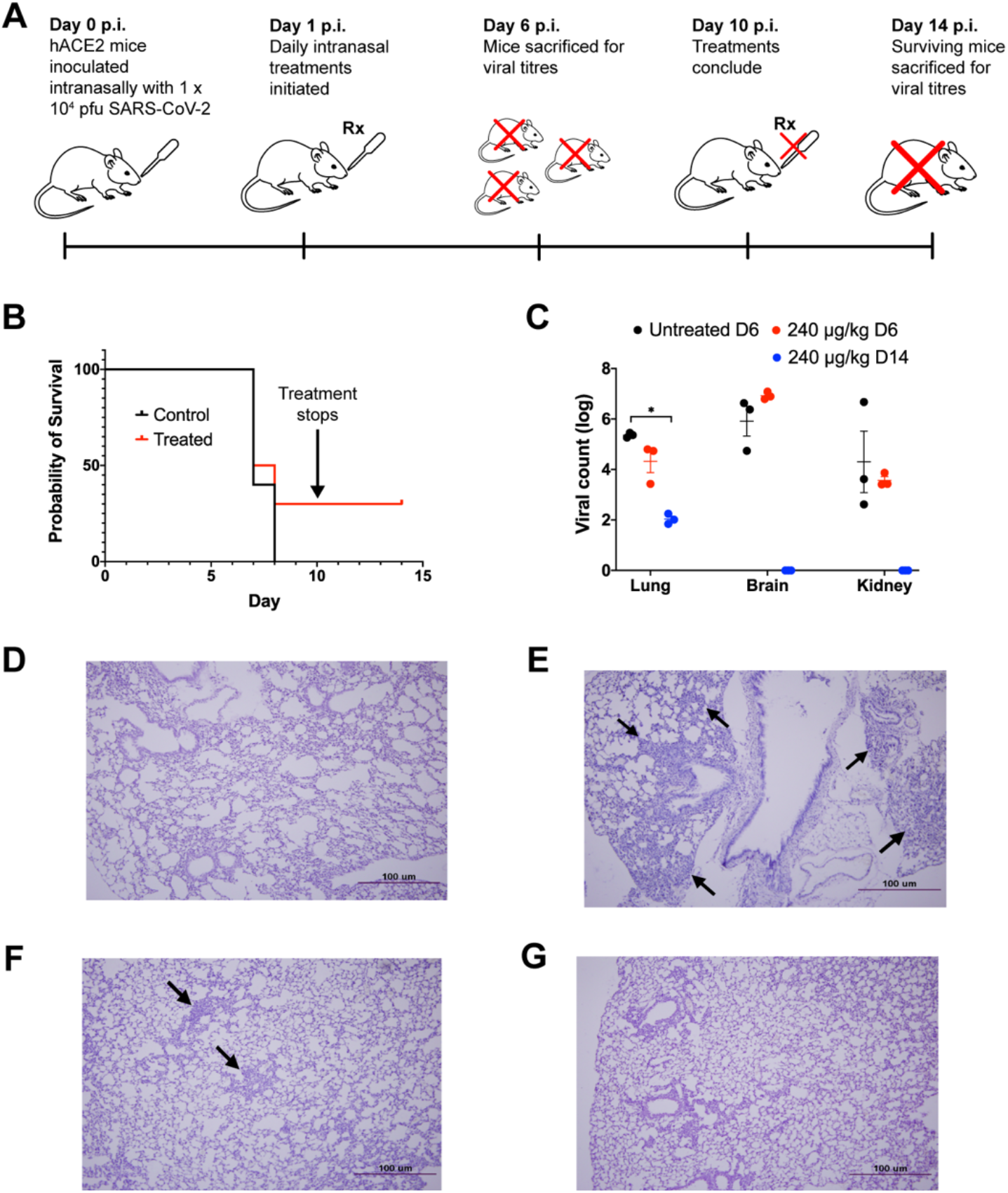
The efficacy of NIC-hLYS particles was assessed in a lethal SARS-CoV-2 infection model (A). Once daily intranasal administration of NIC-hLYS particles suspended in 0.45% NaCl resulted in 30% survival in a lethal SARS-CoV-2 infection (B) and produced a statistically significant decrease in lung viral titers after 10 days of dosing (C). The viral particles obtained from lung and brain homogenates of surviving animals did not produce CPE when administered to Vero E6 cells, indicated that they were no longer active. Compared to lung tissue of uninfected mice (D), infection with SARS-CoV-2 resulted in the development of interstitial pneumonia without treatment (E). By Day 6 of treatment with 240 μg/kg NIC, interstitial pneumonia was notably reduced (F) and further resolved by Day 14 (F). Data are presented as mean + SEM (n = 3). *p < 0.05, using two-way ANOVA with Dunnet’s multiple comparisons test.

In both MERS-CoV and SARS-CoV-2 infection models, the lung tissue of infected and NIC-hLYS-treated mice (Fig. 2F and 3F) showed lower levels of interstitial pneumonia than that of infected and non-treated mice (Fig. 2E and 3E) on day 6 p.i.. Inflammation was further reduced in the treated/infected groups on Day 14 p.i. (Fig. 2F and 3F) and was more comparable to the mock-infected mice (Fig. 2D and 3D), which showed no signs of interstitial pneumonia.

### Niclosamide-lysozyme particles protect against two COVID-19 sequalae: secondary bacterial infection and inflammatory response

COVID-19 patients may be at-risk for secondary bacterial pneumonias and severe inflammatory lung damage. The efficacy of NIC-hLYS in the treatment of these important COVID-19 sequalae was therefore assessed. A resazurin-based microbroth dilution assay was performed to determine the inhibitory activity of several NIC formulations (NIC-hLYS, NIC-BSA, NIC-M, and NIC-DMSO). Compared to the other NIC formulations, NIC-hLYS reached an MIC50 at lower levels of NIC (0.0625 μg/mL), and 100% inhibition was noted at a concentration of 0.125 μg/mL (Fig 4A-C). No inhibitory activity was observed for hLYS alone. Interestingly, all NIC formulations exhibited a similar dose-response profile where a sharp dip in activity preceded the concentrations at which 100% inhibition was achieved. This same pattern was also noted in the dose-response profiles for anti-MERS-CoV and anti-SARS-CoV-2 activity (Fig 1D and E). Plating of the wells with 100% inhibition noted resulted in the growth of colonies at all drug concentrations tested, which indicates that the antimicrobial activity of NIC may be bacteriostatic rather than bactericidal.

**Figure 4:**
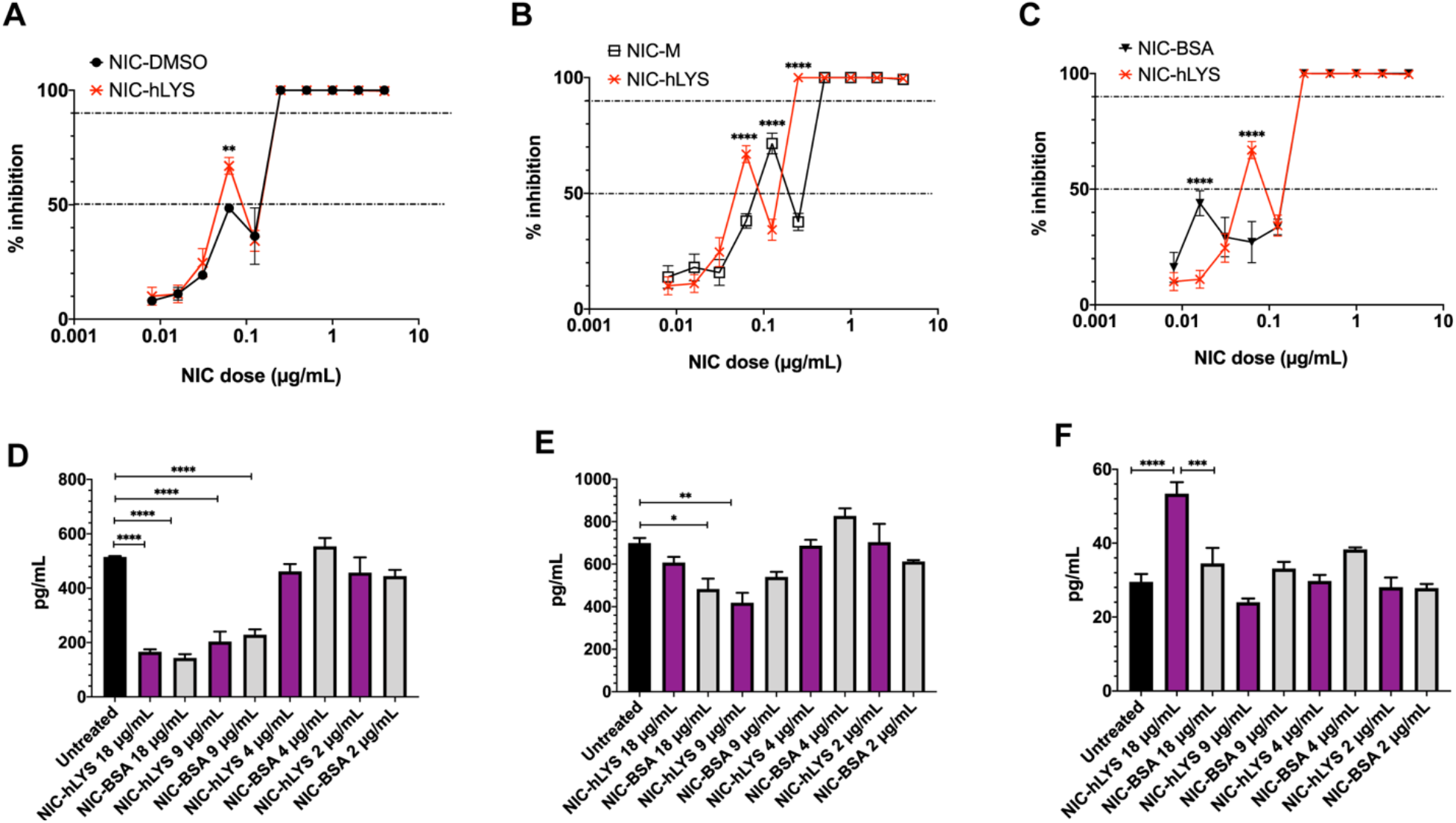
NIC-hLYS resulted in 50% inhibition of the S. aureus Mu50 strain at lower doses than several other NIC formulations tested, included solubilized NIC (A-C). Compared to NIC particles alone, 100% inhibition was achieved at a lower dose (B). Data presented as mean + SEM (n = 6), **p < 0.01,****p < 0.0001, using two-way ANOVA with Dunnet’s multiple comparisons test. NIC-hLYS significantly reduced production of the inflammatory cytokines IL-6 (D) and TNF-α (G) in THP-1 macrophages stimulated with 10 ng/mL lipopolysaccharide (LPS), though a significant increase in IL-1β production was noted at the highest dose tested compared to both the untreated control and NIC-BSA formulation, which may point towards the role of hLYS in inducing production of this cytokine. Data presented as mean + SEM (n =3), *p < 0.05, ** p < 0.01, ***p < 0.001, ****p < 0.0001 using one-way ANOVA with Sidak’s multiple comparisons test.

A feared consequence of SARS-CoV-2 infection is the development of ARDS, which is a major contributor to morbidity and mortality and dramatically increases the burden on healthcare systems. ARDS is caused by the massive release of inflammatory cytokines in the lungs, which occurs in some patients in response to pathogenic infiltration. Both NIC and hLYS are known to exhibit anti-inflammatory activity. The anti-inflammatory activity of two NIC formulations, NIC-BSA and NIC-hLYS, was assessed using an acute macrophage inflammation model. NIC-hLYS and NIC-BSA exhibited similar suppression of the inflammatory cytokines IL-6 and TNF-α (Fig 4D and E). The similarities in the levels of these two cytokines across the two formulations tested indicates that the suppression may be related to the activity of NIC rather than hLYS. Suppression of IL-1β was not observed for either formulation, and the highest concentration of NIC-hLYS tested (18 μg/mL total powder) resulted in statistically significant increase in compared to both the untreated, LPS-stimulated control and an equivalent dose of the NIC-BSA formulation (Fig 4F). Of, note this is an equivalent powder concentration to the dose exhibiting the highest efficacy in the *in vitro* antiviral assays.

### Composite niclosamide-lysozyme particles can be delivered to the upper and lower respiratory tracts with three different commercially available devices

Targeted delivery of antivirals to the respiratory tract carries substantive benefits for the treatment of COVID-19, particularly for compounds with limited oral bioavailability like NIC. However, the wide range of symptoms and disease severity associated with COVID-19 makes development of a broadly applicable therapy difficult. The clinical applicability of our novel NIC-hLYS formulation was assessed by determining the delivery efficiency of the drug using three commercially available respiratory drug delivery platforms: a disposable DPI (TwinCaps®), a vibrating mesh nebulizer (Aerogen Solo®) and a nasal spray (VP7 Aptar®). The composition of spray dried NIC-hLYS particles was optimized using a constrained-mixtures design of experiments (DoE) to achieve respirable dry particles (geometric median diameter ≤ 5 μm) that could be easily reconstituted as suspension suitable for nasal spray or nebulizer-based administration. The DoE generated several promising powder formulations (Supplementary Table 1), of which formulation 8 (F8) was selected for further characterization.

For ambulatory patients, DPIs provide a convenient treatment option for lung-targeted delivery. The rapid administration time for the device as well as the compact size improves patient acceptability and compliance. A disposable DPI is likely to be preferred in the treatment of COVID-19, given the currently unknown risks regarding re-infectability. A commercially available disposable DPI, the TwinCaps® (Hovione) was therefore selected as an exemplary delivery platform. NIC-hLYS powder inhalation was successfully delivered using the TwinCaps DPI, with a 136.0 ± 7.4 μg NIC fine particle dose (i.e., recovered drug mass with less than 5 μm aerodynamic diameter) achieved per 60 mg total powder actuation (0.7% NIC content) when using inhalation flow rate conditions reflective of a healthy patient (Fig 5H). Given the potential for shortness of breath and reduced inspiratory abilities that may occur in COVID-19, the delivery efficiency was also examined using inhalation flow-rate conditions reflective of a patient with reduced lung capacity due to illness or age. Similar fine particle doses (121.1 ± 7.7 μg) were achieved in these reduced inhalation flow rate conditions, which indicates a minimal dependence on inspiratory flow rate to achieve successful delivery of the drug to the peripheral lung regions.

**Figure 5:**
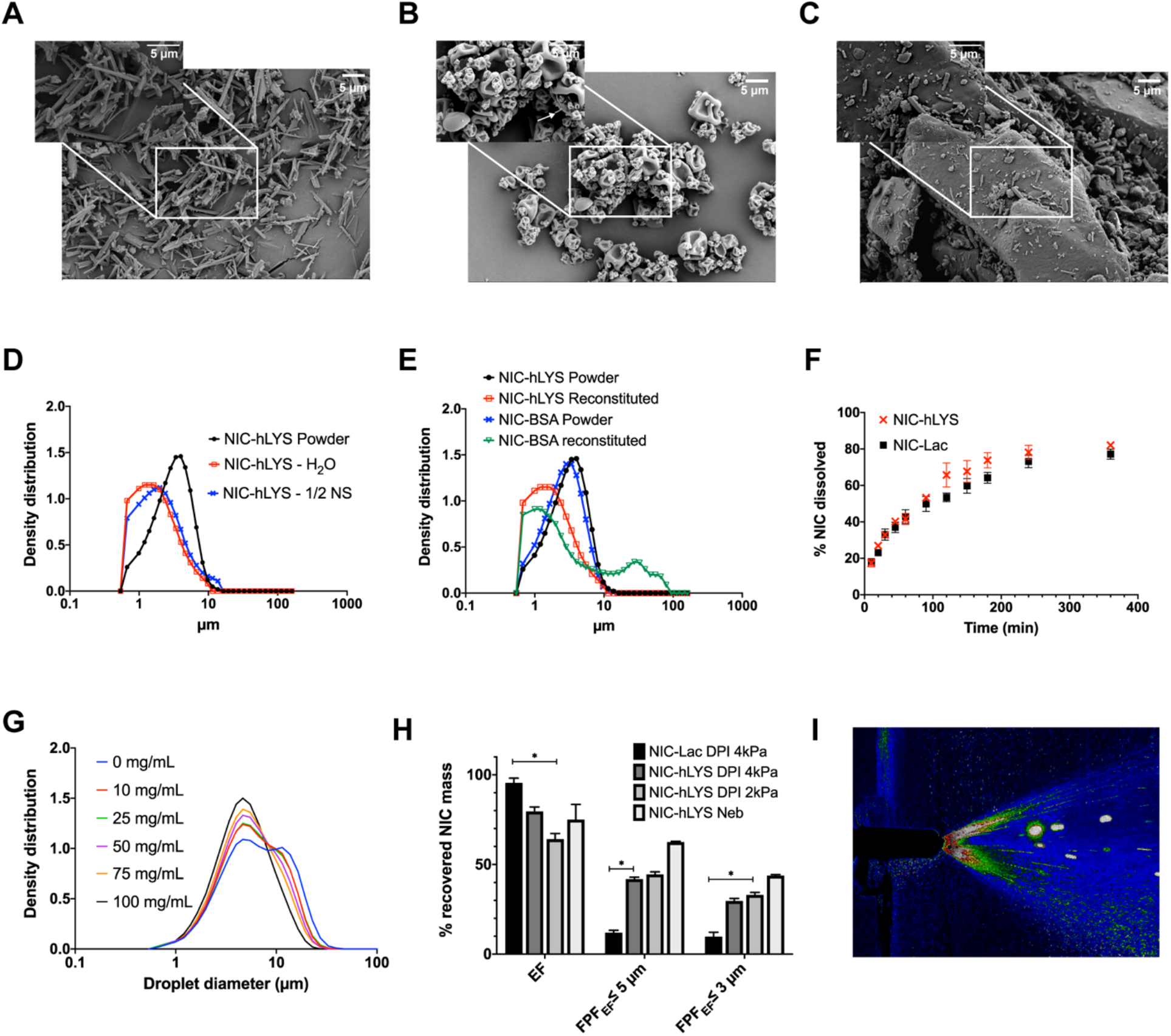
Micronized niclosamide (A) was embedded in a matrix of human lysozyme and stabilizers using spray drying (B). This novel system was developed as an alternative to traditional lactose-based carrier systems (C) and enabled the targeted respiratory delivery of NIC as a powder via DPI or a reconstituted suspension via nebulizer or nasal spray. The optimized formulation exhibited a size distribution that was appropriate for inhalation (i.e., geometric median diameter < 5 μm) in both the dry powder state as well as when reconstituted using water or 0.45% NaCl (D). Similar effects could not be achieved when a negatively charged protein, bovine serum albumin, was substituted in the formulation for the positively charged hLYS (E). Though hLYS is surface active, it appeared to only slightly enhance the dissolution of NIC compared to NIC particles blended with lactose (F). A respirable droplet size distribution could be achieved with multiple different reconstituted concentrations when nebulized using the Aerogen Solo (G). These concentrations resulted in no aggregation to the lysozyme component. Efficient aerosol delivery was achieved with both the nebulizer and disposable DPI, with ~50% of the emitted dose being of an appropriate size for lung deposition. This was significantly improved compared to a traditional lactose carrier particle system (H). Reproducible plume geometry could be achieved using a variety of reconstituted concentrations when actuated using the Aptar device (I). Data is presented as mean + SEM (n = 3). *p < 0.05, using two-way ANOVA with Tukey’s multiple comparisons test (comparisons of DPIs presented only).

COVID-19 may result in the need for mechanical ventilation for continued patient survival. The delivery efficiency of reconstituted NIC-hLYS particles was assessed using an Aerogen Solo vibrating mesh nebulizer, which can be utilized aerosol drug delivery in-line with a ventilator circuit. NIC-hLYS powder reconstituted in 0.45% sodium chloride to a 25 mg/mL concentration (equivalent to 175 μg/mL NIC) resulted in the delivery of a fine particle dose of 62.3 ± 6.4 μg NIC after a 2-minute run time (Fig 5H). Furthermore, a range of concentrations (10 to 100 mg/mL) could be successfully emitted using the Aerogen Solo device (Fig 5G). By altering the reconstitution concentration, the dose of NIC-hLYS could therefore be adjusted if required for pediatric patients, or those with hepatic or renal insufficiencies. The zeta potential of the reconstituted NIC-hLYS powder was determined to be +1.8, in contrast to the poorly performing NIC-BSA particles, which exhibited a zeta potential of −10.9 when reconstituted in water.

Epithelial cells of the upper respiratory tract (i.e., nasal passages) exhibit significantly higher expression of ACE2 receptors than those of the lower respiratory tract, which indicates these tissues may be more prone to infection with SARS-CoV-2(47). As such, the feasibility of administration of the NIC-hLYS formulation using a nasal spray was assessed using spray pattern and plume geometry analysis. NIC-hLYS powders were reconstituted in 0.45% sodium chloride at concentrations ranging from 10 to 50 mg/mL and actuated using a VP7 Aptar® nasal spray device. Suitable plume angles and uniform spray patterns for nasal administration were achieved for all tested concentrations (Fig 5I). An increase in plume angle was noted at increasing powder concentrations which was inversely related to changes in suspension viscosity (Supp Table 2) and may be reflective of the decreased surface tension resulting from the presence of hLYS and other surface-active stabilizers. The utilization of a slightly hypotonic reconstitution medium (Supp Table 3) was selected and used in the *in vivo* studies to improve absorption and potentially assist nose-to-brain penetration of NIC-hLYS. The decreases in brain and kidney viral titers noted *in vivo* when NIC-hLYS was administered intranasally may reflect an achievement of therapeutic drug concentrations outside the respiratory passageways, though this will be evaluated further in a future pharmacokinetic study.

NIC is a poorly water-soluble drug, which renders the commercially available oral formulation ineffective against respiratory diseases due to the limited absorption of the drug from the gastrointestinal tract. Direct delivery to the airways represents a promising alternative to oral delivery, as it would enable achievement of high levels of drug at the site of disease. However, limited solubility and delayed dissolution of niclosamide particles in the upper respiratory tract could result in rapid clearance of the particles by the mucocilliary escalator or through alveolar macrophage uptake. hLYS exhibits surface active properties which could enhance the dissolution rate of poorly water soluble NIC particles. To test this hypothesis, the dissolution rate of NIC-hLYS particles exhibiting an aerodynamic diameter of ~2 μm was compared against hLYS-free NIC particles in simulated lung fluid medium. Though the deposited particles exhibited equivalent sizes and surface areas, the inclusion of hLYS as a formulation component resulted in a slightly faster rate of dissolution, with 82.1% the deposited dose of NIC dissolved by 6 hours (Fig 5F).

### Human lysozyme exhibits stability after processing and aerosolization

Protein aggregation may result in a loss of therapeutic activity or an immunogenic response (48–50). We have previously demonstrated that hen egg white lysozyme is robust to process-induced aggregation using typical particle engineering techniques(51), which provided part of the rationale for its selection as a therapeutically active carrier for the aerosol delivery of NIC particles. Using size exclusion chromatography (SEC), we investigated the formation of higher molecular weight hLYS aggregates before and after spray drying. No further increases in higher molecular weight aggregates were noted after spray drying (Supp Fig 1B) or after nebulization at varying reconstituted concentrations (Supp Fig 1C-F) (Supp Table 4). Interestingly, a decrease in the percentage of solubilized aggregates was noted in the spray dried hLYS formulations compared to the initial unprocessed hLYS product, which may be explained by the shift in secondary structure from towards a higher percentage of parallel β-sheet structure upon spray drying, whereas the unprocessed hLYS had a higher percentage of anti-parallel β-sheet structure (Supp Table 5). The glass transition temperature (Tg) measured for the NIC-hLYS powder was 79.4°C, which makes it suitable for storage in ambient conditions without risk of further protein degradation. The water content of the spray dried powder was determined to be 8.8% based upon Karl Fisher coulometric titration, which is similar to literature reported values for the water content of lysozyme.(52)

## DISCUSSION

The co-formulation of the cationic, endogenous protein hLYS with micronized NIC produced a 4-fold increase in potency against CoVs and a 2-fold increase in potency against MRSA compared to NIC particles alone. Though the inclusion of hLYS did slightly increase the dissolution rate of micronized NIC particles, this alone cannot explain the increased potency, as solubilized NIC exhibited lower antiviral activity at the 0.125 μg/mL dose than both NIC-M and NIC-hLYS. hLYS plays an important role in the innate immune system, and it is found in abundance at the mucosal surfaces of the respiratory tract as well as in the lysosomal granules of neutrophils and macrophages.(53) The antibacterial efficacy of hLYS stems from both its enzymatic activity, which results in the hydrolysis of the glycosidic bonds linking peptidoglycan monomers in bacterial cells walls, as well as its cationic activity, which enables insertion of the protein and formation of pores in negatively charged bacterial membranes.(46) To our knowledge, the activity of lysozyme against coronaviruses has not been previously reported. A peptide (HL9) located within the helix-loop-helix motif of hLYS has been previously demonstrated to block HIV-1 viral infection and replication, whereas mutants of HL9 did not.(43) The location of this peptide was deemed to be separate from the hydrolytic site. It is possible that this peptide sequence, or another, is responsible for the anti-coronavirus activity of hLYS.

Cationic peptides have been hypothesized to exhibit antiviral activity due to a disruption of the viral particle membrane(54). hLYS may also disrupt various signaling pathways (TGFβ, p53, NFκB, protein kinase C, and hedgehog signaling) which affect host cell susceptibility to viral infection.(43) An immunomodulatory response may explain the notable delay in anti-SARS-CoV-2 activity at lower doses when hLYS was used alone. Additionally, a unique up-regulation in IL-1β activity was noted for the NIC-hLYS formulation in a macrophage model of inflammatory stimulation, while expression of other inflammatory cytokines (IL-6, TNF-α) was decreased. IL-1 family cytokines have been previously associated with the induction of antiviral transcriptional responses in fibroblasts and epithelial cells.(55) In SARS-CoV infected African green monkeys, significantly lower levels of IL-1β were noted in the lungs of aged monkeys compared to juvenile monkeys.(56) Similarly, elderly mice infected with influenza exhibited lower levels of 1L-1β, and administration of IL-1β augmenting compound improved morbidity and mortality.(57) Thus, it is possible that a similar immune-mediated mechanism is contributing to the increased activity of NIC-hLYS compared to other NIC formulations, though this requires further investigation.

At physiological pH, hLYS exhibits a positive charge while NIC exhibits a negative charge. This attraction may contribute to the improved dispersibility of NIC suspensions when hLYS is utilized as the carrier protein, as demonstrated by the notable aggregation observed when a negatively charged protein, BSA, is used in the formulation. This charge-based interaction may have additional benefits when NIC is in the solubilized state. NIC has two substituted aromatic rings, the electronegativity of which has been deemed critical for its activity against other viruses(9). Non-covalent interactions between aromatic compounds is a known phenomenon, which occurs as a result of the overlap of the π-orbital of the two electron clouds of the aromatic rings(58, 59) and π-π stacking of NIC is observed in the crystalline form.(60) Though not investigated in this study, it’s possible that self-association of NIC molecules may disrupt the availability of the strongly electro-negative groups necessary for antiviral activity. This may explain why a decrease in antiviral activity was noted with NIC solubilized in DMSO compared to NIC particles. Interactions between NIC and hLYS in the microenvironment may serve as a mechanism to disrupt this self-association and ensure availability of the electronegative functional groups, as lysozyme has been previously demonstrated to complex with negatively charged molecules (61) and a hydrophobic drug (62). Given the physicochemical properties of NIC, both charged-based and hydrophobic interactions with hLYS may be important. Future studies will examine the interactions and mechanisms of complexation between NIC and hLYS to further elucidate how the molecular interactions may contribute to improved antiviral activity.

At present, there are limited treatment options for COVID-19. In August 2020, the FDA approved remdesivir for the treatment of all hospitalized adult and pediatric patients with COVID-19, irrespective of the severity of disease.(63) This approval is based upon a statistically significant reduction in median time to recovery for patients treated with remdesivir in a recent clinical trial.(64) However, remdesivir is currently not recommended for use in patients with acute or chronic kidney disease (GFR < 30 mL/min), which may limit its utility in severe COVID-19 infection. This contraindication stems from the incorporation of sulfobutylether-β-cyclodextrin (SBECD), which is utilized as a vehicle for the poorly water soluble remdesivir in the intravenous product, and which can accumulate in cases of renal failure. Clinical trials examining inhalation-based delivery of remdesivir are currently under way(65, 66), though details of this aerosol formulation and whether or not it contains SBECD are not provided. A pre-print of a powder aerosol formulation of remdesivir has also been recently published(67). These studies overall indicate the clinical interest in the development of an inhalation-based treatment for COVID-19.

Based upon our data, NIC may be a promising alternative or adjunt therapy to remdesivir for the treatment of COVID-19. A previous study examining the efficacy of remdesivir for the treatment of a lethal MERS-CoV infection (5 × 10^5^ pfu) followed by treatment 24 hours p.i with remdesivir resulted in ~2.5-3 log reduction in viral particles in the lung tissue(68), which is similar to the reduction noted in our study using intranasal NIC-hLYS. At the day 6 p.i. endpoint of the remdesivir MERS-CoV study equivalent survival (50%) was noted in the treated and untreated groups. In comparison, at day 7 p.i., we noted 71.4% survival in treated animals versus 60% in untreated controls, and at day 10 p.i. 43% of treated animals had survived while 0% of untreated animals had survived. Notably, these animals continued to survive after treatment was ceased, which was reproduced in the lethal SARS-CoV-2 infection model. While it is difficult to make comparisons between the two drugs on survival improvement given differences in study length and inoculum concentration, it appears survival with NIC-hLYS treatment is comparable if not improved compared to remdesivir treatment in lethal MERS-CoV infections. Expansion of the dosing range or frequency in future studies, as well as the incorporation of prophylactic dosing may enable a significant improvement in survival to be achieved, similar to what was noted with remdesivir when used prophylactically *in vivo*. Furthermore, we have demonstrated *in vitro* efficacy of NIC-hLYS reducing complications related to COVID-19, most notably secondary bacterial pneumonia by MRSA and dampening (but not complete suppression) of TNF-α and IL-6, which are known markers for COVID-19 disease severity(69). This broad-spectrum activity may provide a unique advantage for NIC-hLYS compared to other leading COVID-19 candidates as an early report on COVID-19 reported 50% of patients that died had a secondary infection(70). Though NIC appears to be primarily renally cleared(71), limited data is available regarding the effects that hepatic or kidney failure may have on the toxicity profile of the drug. Future studies will evaluate the pharmacokinetic distribution of NIC-hLYS delivered via the intranasal or pulmonary routes.

We have successfully demonstrated NIC-hLYS is effective *in vitro* and *in vivo* for the treatment of COVID-19, and importantly, scalable inhaled drug delivery systems can be developed based upon the formulation which will enable rapid availability to a global patient population. Historically, it has been observed that variations in patient inspiratory force can have profound effects on the magnitude and region of dose delivery to the airways from DPIs.(72) One of the most common symptoms of COVID-19 is shortness of breath, reportedly occurring in 50.8% of patients.(73) Moreover, patients with severe infection were significantly more likely to have shortness of breath than patients with non-severe infection.(73) NIC-hLYS powder exhibited efficient aerosol performance using a disposable DPI device even at reduce inspiratory efforts which will ensure reproducible dose delivery throughout disease progression, as well as in pediatric patients. The powder can be reconstituted at the point of care to form a stable suspension appropriate for nebulization, thus enabling treatment of both ambulatory and severely ill or mechanically ventilated patients. Lastly, NIC-hLYS suspension could be reproducibly actuated using a commercially available nasal spray device and has demonstrated promising antiviral activity *in vivo* when administered to infected mice via the intranasal route. The utilization of a slightly hypotonic carrier may improve systemic penetration from the nasal route and promote distribution in the brain(74, 75). SARS-CoV-2 has been found to exhibit neuroinvasive properties, and severely affected COVID-19 patients appear to be more likely to develop neurological symptoms compared to those with mild disease(76, 77). We observed that the brain tissue of surviving, NIC-hLYS treated animals with SARS-CoV-2 infection exhibited no detectable viral particles. NIC-hLYS delivered by the intranasal route may therefore be effective in preventing viral invasion of brain tissue or reducing viral replication within brain tissue.

We acknowledge limitations within our study, namely the lack of pharmacokinetic data which may explain the mechanism of viral load reduction in brain, kidney, and lung tissue, as well as the utilization of only one dosing level of NIC-hLYS in the SARS-CoV-2 model. In our study, we utilized a “worst-case scenario” of efficacy evaluation by administering lethal inoculums of CoVs and initiating treatment 24 hours after the infection was established(78). The effect of the timing of treatment initiation, i.e., prophylactic versus 24 or 48 hours p.i., was not examined. Incorporation of prophylactic dosing may result in the increased efficacy of NIC-hLYS, as the antiviral mechanism of action of NIC may be related to the prevention of viral entry into host cells.(9) Future studies will examine these variables, based upon the notable activity of the novel NIC-hLYS formulation in these initial proof-of-concept efficacy studies.

In conclusion, a novel formulation of NIC-hLYS optimized for delivery to the upper and lower respiratory tracts as a powder or stable suspension was developed. *In vitro*, the incorporation of hLYS into the formulation was noted to improve potency against MERS-CoV and SARS-CoV-2, as well as MRSA, which is an important causative agent for secondary bacterial pneumonias associated with COVID-19. The NIC-hLYS particles exhibited suppression of the inflammatory cytokines TNF-α and IL-6, which have been implicated in the development of more severe COVID-19, while elevating the production of the inflammatory cytokine IL-1β, which may contribute to enhanced antiviral activity. Intranasal administration of NIC-hLYS particles improved survival and reduced viral tissue loads *in vivo* in two lethal CoV infections at a level that was comparable to the leading COVID-19 treatment candidate, remdesivir. Thus, NIC-hLYS is likely to not only have efficacy in the treatment of the current SARS-CoV-2 pandemic but could be utilized as a treatment in future CoV pandemics.

## METHODS

### Engineering and optimization of inhalable composite particles of niclosamide and human lysozyme

Niclosamide (NIC) was obtained from Shenzhen Neconn Pharmtechs Ltd. (Shenzhen, China) and micronized in-house using an Model 00 Jet-O-Mizer air jet mill (Fluid Energy Processing and Equipment Co, Telford, PA, USA) using a grind pressure of 75 PSI and a feed pressure of 65 PSI for a total of three milling cycles to achieve an x50 diameter of 2.2 μm and an x90 diameter of 4.1 μm. To generate a powder formulation of NIC suitable for DPI-based delivery as well as suspension-based nebulizer and nasal spray delivery, micronized NIC particles were embedded in a matrix of recombinant human lysozyme (hLYS) (InVitria, Junction City, KS, USA), sucrose (Sigma-Aldrich, Darmstadt, Germany), polysorbate 80 (Sigma-Aldrich) and histidine (Sigma-Aldrich) using spray drying. Our group has previously identified that histidine (buffering agent), sucrose (lyoprotectant agent), and polysorbate 80 (surface active agent) can be used to generate stable and dispersible formulations of lysozyme for delivery via DPI (51, 79).

Preliminary screening experiments indicated that spray drying with a feed solid content greater than 1% w/v resulted in a dry particle size distribution (PSD) that was greater than the respirable size (typically less than 5 μm particle diameter). A constrained mixtures DoE was therefore utilized to determine the optimal ratio of micronized niclosamide, human lysozyme, sucrose, and polysorbate 80 in the 1% w/v feed to generate powder with both a dry and reconstituted PSD suitable for oral or nasal inhalation (Supplementary Table 6).

The upper constraint of polysorbate 80 was selected to match the maximum FDA approved level of the excipient for the inhalation route.(34) The lower level of human lysozyme was set at 60% w/w, based upon previously published results indicating that a respirable powder cannot be generated below this level(80). Sucrose and polysorbate 80 lower level constraints were set at conservative values, as their inclusion is necessary to ensure stability of lysozyme during the drying process, but the lowest level needed for stability has not been defined. The micronized niclosamide lower and upper constraints were selected to be around the point at which the average number of particles is excepted to be one, based upon equation 1(81).

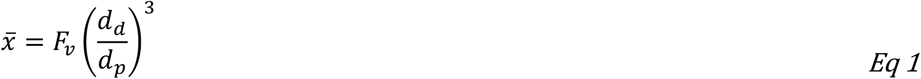

Where 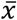 is the average number of particles per droplet, *F*_*v*_ is the volume fraction of particles of diameter *d*_*p*_, *d*_*d*_ is the diameter of the particles (assumed to be equivalent to the X90 diameter of the micronized niclosamide particles) and *d*_*p*_ is the diameter of the droplets (assumed to be equivalent to the X90 diameter of the atomized droplets at conditions used in the experiment).

A constrained mixtures DoE was generated using the RSM package(82) in the opensource software R(83). Of these 25 mixtures, a 12-run *D*-optimal subset (Supplementary Table 7) was generated and prepared using spray drying to enable fitting of the results to the Scheffé quadratic model. The dry components of the formulations were mixed using a process of geometric dilution and wetted and suspended using polysorbate 80 followed by incremental additions of 0.174 mg/mL histidine buffer. All suspensions were spray dried with a BUCHI B-290 mini-spray dryer (BUCHI Corporation, New Castle, DE, USA) coupled to a syringe pump (KD Scientific Inc, Holliston, MA, USA) set at a feed rate of 1 mL/min. A 2-fluid pneumatic atomizer nozzle (0.7 mm with 1.5 mm cap) was used to atomize the suspension, and house air was used as the atomization gas. The cleaning needle of the nozzle was removed to prevent disruptions to the feed flow rate.(51) The spray dryer was set at an inlet temperature of 130°C, which corresponded to an outlet temperature of ~70°C. For all runs, no settling of the feed suspensions was noted during processing. Formulations were evaluated on the basis of dry powder PSD and reconstituted suspension PSD, and the composition exhibiting the most promising characteristics was selected for further evaluation. Comparative powders were generated for the purposes of evaluation of physicochemical characteristics, aerosol performance, and efficacy. A NIC-free hLYS spray dried powder was generated using the optimized formulation composition identified in the DoE, minus the addition of micronized NIC. To compare the novel NIC-hLYS powders against a traditional lactose-based carrier system, micronized NIC was blended with crystalline lactose particles (Lactohale 100; DFE Pharma) using geometric dilution followed by mixing in a Turbula powder blender. The concentration NIC in the niclosamide-lactose blend (NIC-Lac) was set to match that in the NIC-hLYS powder (0.7%). Lastly, a powder was generated in which bovine serum albumin (BSA) was substituted for the hLYS in the optimized NIC formulation, in order to assess the effect of the protein on formulation characteristics.

### 2. Physicochemical characterization of niclosamide-lysozyme composite particles

Particle size distribution (PSD) of NIC-hLYS powders was measured using a RODOS disperser coupled to a Sympatec laser diffractor unit (Sympatec GmbH, Clausthal-Zellerfeld, Germany). Dispersion pressure was set at 3.0 bar and feed table rotation was set at 20%. Time slices of the plume exhibiting an optical concentration between 5-25% were averaged to generate the PSD. The PSD of the reconstituted powders was determined in both ½ normal saline (NS) and DI water using the Cuvette attachment for the laser diffraction. A spin bar was set to rotate at 2000 RPM, and the dry powders were added directly to the solvent in the cuvette until an optical concentration exceeding 5% was reached. Three measurements were taken and averaged. Zeta potential of NIC-hLYS suspensions before and after spray drying was determined using a Zetasizer NanoZS (Malvern Panalytical Ltd, Malvern, UK) and compared against a NIC-BSA suspension.

The morphology of NIC-hLYS powders was observed using scanning electron microscopy (SEM). Samples were mounted onto aluminum stubs using double-side carbon tape and sputter coated with 15 nm of platinum/palladium (Pt/Pd) under argon using a Cressington sputter coater 208 HR (Cressington Scientific Instruments Ltd, Watford, UK). Images were obtained using a Zeiss Supra 40VP SEM (Carl Zeiss Microscopy GmbH, Jena, Germany).

Glass transition temperature (Tg) and crystallinity of NIC-hLYS powder was determined using modulated DSC. Powder samples were loaded into Tzero pans with hermetically sealed lids, and a hole was pierced to prevent pan deformation. A scan was performed on a Q20 DSC (TA Instruments, New Castle, DE, USA) by ramping 10°C/min to −40°C, followed by a 2°C/min ramp from −40°C to 280°C with a modulation cycle of ±0.5°C every 40 seconds. Data was processed using TA Universal Analysis.

The dissolution profile of the aerosolized NIC-hLYS and NIC-Lac powders was evaluated based upon previously published methods (84, 85) using a composition of simulated lung fluid (SLF) adapted from Hassoun et al (86). Briefly, Whatman GF/C glass microfiber filters (diameter = 24 mm) were placed in a Stage 4 of a Next Generation Impactor, which corresponded to an aerodynamic size cut off of 2.01 μm at the 40 L/min flow rate used in the experiment. Powders were dispersed using a disposable TwinCaps (Hovione) DPI. The filters were transferred to a modified Transwell system (membrane removed) to enable contact of the bottom of the filter with a basal compartment containing 1.5 mL SLF. The apical side of the filters were wetted with 0.1 mL of SLF. The dissolution results are presented in Table 3. The Transwell system was placed in a 37°C isothermal chamber and 0.1 mL samples were removed from the basal compartment at various timepoints and replaced with fresh SLF.

### 3. Stability analysis of human lysozyme

Effects of processing and nebulization on the aggregation of hLYS was assessed using size exclusion chromatography (SEC) based upon a previously published method (80). The effect of processing on the secondary structure of hLYS was determined using a Niclolet iS50 Fourier transform infrared spectrophotometer with attenuated total reflectance (FTIR-ATR) (Thermo Scientific). Spectra were acquired using OMNIC software from a wavelength of 700 cm^−1^ to 4000 cm^−1^ with 64 acquisitions in total. An atmospheric background scan was collected and subtracted from all powder spectra. Secondary structure analysis was performed in OriginPro (OrginLab Corporation) using the second derivative of the amide I band region (1580-1720 nm) and the peak analyzer function. The region of interest was first baseline-corrected, and the second derivative of the spectra was smoothed using the Savitzky-Golay method with a polynomial order of 2 and 50 points in the smoothing window. Peaks identified from the second derivative minimums were iteratively removed to assess the effect on the model fit.

### 4. Aerosol performance testing

The aerosol performance of the spray-dried composite NIC-hLYS powder was assessed using a disposable TwinCaps DPI. Performance was assessed at both a 4kPa and 2kPa pressure drop through the device to determine the effects of inspiratory flow rate on emitted and fine particle dose. For comparative purposes, the performance of traditional lactose carrier-based dry powder, NIC-Lac, was also assessed. A 4 kPa pressure drop was generated through the TwinCaps DPI using an inspiratory flow rate of 40 L/min. A 6.5 second actuation time was used to pull 4 L of air through the NGI. A 2 kPa pressure drop was generated using an inspiratory flow rate of 28.3 L/min, and an 8 second actuation time was used. For all experiments, 60 mg of powder was loaded into the device. After actuation of NIC-hLYS powders, niclosamide was collected by dissolving the deposited powder using a 50-50 water:acetonitrile mix. An aliquot was taken, and an additional volume of acetonitrile was added to bring the final ratio to 20:80 water:acetonitrile. To induce phase separation, A 2M solution of ammonium acetate was added to this mixture at a volume that was 20% of the water:acetonitrile mix. Niclosamide was assayed from the upper organic layer by measuring absorbance at 331nm using a plate reader. For the niclosamide-lactose blend, the deposited powder was collected by dissolving it in 20:80 water:acetonitrile, centrifuging, and then measuring absorbance at 331nm.

Delivery of the reconstitued NIC-hLYS suspension was assessed using the disposable Aerogen® Solo vibrating mesh nebulizer (Aerogen). Preliminary screening experiments indicated that a reconstituting 25 mg/mL of NIC-hLYS powder in 0.45% w/v sodium chloride (1/2 NS) reduced the changes in nebulizer concentration during therapy compared to higher concentrations; therefore, this concentration was utilized for further analysis. The inspiratory flow rate was set for 15 L/min and the apparatus was chilled to 4°C as specified by the United States Pharmacopeia (USP)(87). Nebulization was performed for two minutes to ensure sufficient deposition of drug in the stages for analysis.

After drug collection, aerosol performance was evaluated on the basis of emitted fraction or dose (niclosamide mass emitted from the device as a percentage of the total recovered powder) and fine particle fraction (niclosamide mass with a size cut off of less than 5 μm aerodynamic diameter or 3 μm aerodynamic diameter, as a percentage of the emitted dose). NGI stage cut-offs were determined for the flow rate utilized based upon Eq. 2, while the MOC cut-off diameter was determined using Eq. 3.

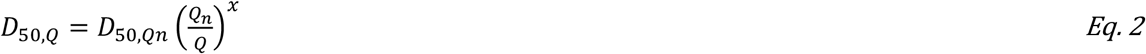

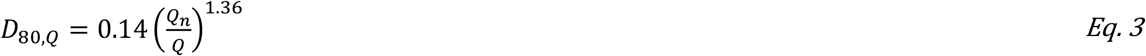

Where *D*_*50,Q*_ is the cut-off diameter at the flow rate *Q*, *D*_*50,Qn*_ is the cut-off diameter at the archival reference values of *Q*_*n*_ = 60 L/min, and the values for the exponent, *x*, are those obtained from the archival NGI stage cut size-flow rate calculations determined by Marple et al(88). Aerosol performance was evaluated on the basis of emitted fraction (EF), which is defined as the cumulative mass emitted from the device as a fraction of the recovered mass, fine particle fraction less than 5 μm (FPF_<5μm_), defined as the mass less than 5 μm aerodynamic diameter as a fraction of the emitted dose, and the fine particle fraction less than 3 μm (FPF_<3μm_) The FPF_<5μm_ and FPF_<3μm_ values were interpolated from a graph plotting the cumulative percentage of NIC deposited in a stage against the cut-off values of the stage.

### 5. Nasal spray characterization

To evaluate the utility of the optimized NIC-hLYS powders for nasal administration, suspensions of varying concentrations (10 mg/mL, 25 mg/mL, and 50 mg/mL) were prepared in 1/2 NS and placed in a VP7 pump Aptar® pump meter spray device. Spray patterns and plume geometries were evaluated using laser-assisted high speed imaging based on methods previously reported by Warnken et al.(89) Briefly, the loaded spray devices were actuated using a MightyRunt automated actuator (InnovaSystems, Inc) set at parameters that mimic those of an average adult user(90). A laser-sheet was oriented either parallel or perpendicular to the actuated spray at distances of 2 and 5 cm from the nozzle tip in order to assess the plume geometry and spray pattern, respectively. The actuation was conducted in a light-free environment in order to isolate the portions of the spray photographed by the high-speed camera (Thorlabs, Inc.) from those illuminated in the plane of the laser. Image analysis of the plume geometry and spray pattern were permed in Fiji.(91) For the plume geometry, the outline of the observed plume was traced and the slope of each side of the plume was determined. This was used to calculate the angle formed at the intersection of the two lines. The spray pattern characteristics including maximum and minimum diameters were determined using the software’s measurement function to determine the Feret diameters.

### 6. *In vitro* anti-inflammatory efficacy

The ability of the optimized NIC-hLYS formulation to dampen inflammatory response was evaluated using lipopolysaccharide (LPS) stimulation in a THP-1 macrophage model. THP-1 monocytes were seeded in 6-well plates at a concentration of 4 × 10^5^ cells/mL (5 mL total) in RPMI 1640 media supplemented with 10% FBS, 1% penicillin/streptomycin, and 15 ng/mL phorbol 12-myristate (PMA) to induce differentiation into mature macrophages. The cells were incubated in the presence of PMA for 48 hours, after the media was replaced with PMA-free media and cells were rested for 24 hours. NIC-hLYS and NIC-BSA powders were suspended in RMPI 1640 media at varying concentrations (2 to 18 μg/mL, based on total powder content) and added to the cells simultaneously with 10 ng/mL LPS. The cells were then incubated for 6 hours to achieve peak cytokine expression(92). Following incubation, supernatants were collected, and cytokine concentrations were quantified using ELISA (DuoSet, R&D Systems) and compared against untreated controls.

### 7. *In vitro* efficacy

All antiviral efficacy experiments were performed using Vero-E6 cells obtained from American Type Culture Collection (Manassas, Virginia, USA). Vero-E6 cells were maintained in Minimal Essential Medium (MEM) supplemented with 10% fetal bovine serum (FBS) and 1X antibiotic-antimycotic solution (Sigma, St. Louis, USA) (i.e., MEM complete). Cells were infected with either MERS-CoV (EMC2012 strain) or SARS-CoV-2 (SARS-CoV-2/human/Korea/CNUHV03/2020 strain). All experimental procedures involving potential contact with MERS-CoV or SARS-CoV-2 were conducted in a biosafety level 3 laboratory of Chungnam National University, which was certified by the Korean government.

#### 7.1. Efficacy determined by viral titer

Vero-E6 cells (2 X10^4^/ml) were seeded in the wells of 6-well tissue culture plates. After a 3-day incubation period, cells were washed with warm PBS (pH 7.4) twice and were infected with SARS-CoV-2 (1.7 × 10^3^ pfu) or MERS-CoV (2 × 10^4^ pfu) diluted in MEM with 2% FBS, which was followed by a 24-hour rest period. The media was then replaced with MEM complete containing various concentrations of the investigational formulations (prepared as suspensions). For the assessment of solubilized NIC, stock solutions were prepared by dissolving the drug in DMSO, and the diluting in MEM-complete, with the resulting media not containing more than 1% DMSO. Drug concentrations and formulations were assessed in triplicate. For each 6-well plate, 1 well was utilized as untreated control. Cells were incubated for 24 or 48 hours, at which point viral RNA from samples was isolated using RNeasy Mini Kit (QIAGEN, Hilden, Germany). Viral RNA was quantified with TaqMan real time fluorescent PCR (RTqPCR) using a TOPreal™ One-step RT qPCR Kit (Enzynomics, Daejeon, Korea) and SARS-CoV-2 and MERS-CoV primers and probe (Supplementary Table 8). Real-time amplification was performed using a Rotor-Gene 6000 (QIAGEN, Hilden, Germany). An initial incubation was performed at 50°C for 30 minutes and at 95°C for 10 minutes, after which 45 cycles of a 5 second hold at 95°C and a 30 second hold at 60°C were performed. Cycle threshold (Ct) values were converted to plaque forming units (pfu) using a standard curve generated from data using stock viruses with known pfu titers by plaque assay.

#### 7.2. EC_50_ determination

Vero-E6 cells grown in tissue culture flasks were detached by treatment with trypsin-EDTA and were seeded in 96-well tissue culture plates. When confluent, cells were washed with warm PBS (pH 7.4) and infected with MERS-CoV or SARS-CoV-2. The half maximal effective concentration (EC_50_) of the formulations was assessed by dosing infected Vero E6 cells plated in 96-wells with NIC-hLYS suspensions with NIC content ranging from 0.25 μg/mL to 0.004 μg/mL once daily over the course of 72 hours. Cell viability was determined on day 4 by observing cytopathic effects (CPE) under microscope. The EC50 was calculated as the concentration of NIC resulting in no observable CPE in 50% of the wells. For comparative purposes, the EC50 of micronized NIC without the inclusion of hLYS was also evaluated.

#### 7.3. Inhibitory activity against S. aureus

The inhibitory activity of various NIC formulations (NIC-hLYS, NIC-M, NIC-BSA, and NIC-DMSO) against methicillin-resistant S. aureus strain Mu50 was assessed using a resazurin-based 96-well plate microdilution assay(93). Varying concentrations of the NIC formulations (: 4, 2, 1, 0.5, 0.25, 0.125, 0.0625, 0.0313, 0.0156, 0.008 μg/mL, dosed based on NIC content) were plated with a 5 × 10^5^ cfu/mL inoculum of *S. aureus* (n = 6 per dose). One column of the plate was used as growth control, i.e., no antibiotics were added, while another column was used as a sterile control, i.e., no bacteria added. The plates were incubated for 24 hours at 37°C with 150 RPM shaking, after which point 30 μL a 0.015% resazurin sodium solution was added. The plates were incubated for an additional 2 hours to allow color change to occur, and the fluorescence of the wells was read at 530 nm excitation/590 nm emission. The fluorescence of the sterile wells was subtracted from the fluorescence of all treated wells, and a decrease in fluorescence of the treated wells versus the growth control was noted as inhibitory activity. The content of wells exhibiting 100% inhibition were plated on tryptic soy agar plates and incubated overnight to determine the mean bactericidal concentration (MBC).

### 8. *In vivo* efficacy assessment

#### 8.1. MERS-CoV infection

HDPP-4 transgenic mice were kindly provided by Dr. Paul B. McCray Jr (University of Iowa). MERS-CoV infection was initiated in anaesthetized mice by intranasal (i.n.) administration of 50 μL (1 × 10^5^ pfu) of MERS-CoV (EMC2012 strain), which was kindly Drs Bart Haagmans and Ron Fouchier (Erasmus Medical Center). Efficacy was initially established using a dose-finding study, in which treatments were initiated 1-day post-infection (p.i.) and daily for 6 days, at which point animals (n = 3 from each group) were sacrificed. NIC-hLYS powder was reconstituted in 0.45% sodium chloride to achieve a dose of 60 or 120 μg/kg NIC (n = 7 per group). The suspensions were administered i.n. in a volume of 50 μL, and 50 μL of 0.9% sodium chloride was administered as a control (n = 6). Though a second timepoint was intended for day 9, death due to illness or as a result of treatment administration prevented obtainment of these data. A separate study was conducted to compare the survival of MERS-CoV-infected mice treated with 240 μg/kg NIC-hLYS (n = 7) and placebo (n = 6). In this study, mice were dosed intranasally for 10 days, at which point treatment was terminated. Surviving mice were rested without treatment for an additional 3 days, and sacrifice was performed Day 14 p.i. to obtain tissues (lung and brain) for viral titres and tissue pathology. The weight of mice was recorded daily. Tissues (0.1 g per sample) were homogenised using a BeadBlaster homogeniser (Benchmark Scientific, Edison, New Jersey, USA) in 1 mL of PBS (pH 7.4) to measure virus titres by RT-qPCR. The remaining portions of tissues were used for histopathology. Mice were lightly anaesthetized with isoflurane USP (Gujarat, India) prior to all viral inoculation and dosing procedures.

#### 8.2. SARS-CoV-2 infection

HACE-2 transgenic mice (K18-hACE2 mice) (The Jackson Laboratory, USA) were lightly anaesthetized with isoflurane USP (Gujarat, India) and inoculated intranasally (i.n.) with 50 μL (1 × 10^4^ pfu) of SARS-CoV-2/human/Korea/CNUHV03/2020. Animals were rested for 24-hours, after which daily treatment was initiated with i.n. NIC-hLYS reconstituted in 0.45% sodium chloride (240 μg/kg NIC) (n = 13) or 0.9% sodium chloride as a placebo (n = 8). All treatments were performed on anaesthetized mice. On day 6 post-infection, 3 mice per group were euthanized, and lung, brain and kidney tissues were collected for viral titres and tissue pathology. Treamtne was performed until 10 days p.i., at which point surviving animals were left untreated for 3 days, and then sacrificed on day 14 p.i. to obtain tissues for viral titres and pathology. Tissues (0.1 g per sample) were homogenised using a BeadBlaster homogeniser (Benchmark Scientific, Edison, New Jersey, USA) in 1 mL of PBS (pH 7.4) to measure virus titres by RT-qPCR. The remaining portions of tissues were used for histopathology.

#### 8.3. Preparation of tissues for histopathology

Mouse tissues were fixed in 10% neutral buffered formalin (10%) and then embedded in paraffin. The lung tissue was cut into 5 μm sections, which were stained with haematoxylin (H) solution for 4 min. The stained tissue sections were washed with tap water for 10 min and then stained with eosin (E) solution. The stained sections were visualised under an Olympus DP70 microscope and photographed (Olympus Corporation, Tokyo, Japan).

#### 8.4. TCID_50_ assay

To determine whether measured viral particles in lung and brain tissue were dead or alive, the log_10_TCID_50_/mL was determined. Vero-E6 cells grown in tissue culture flasks were detached by treatment with trypsin-EDTA and were seeded in 96-well tissue culture plates with MEM containing 10% FBS and 1× antibiotic-antimycotic solution. When confluent, the cells were washed with warm PBS (pH 7.4) and infected with virus samples, which were 10-fold diluted in MEM with 2% FBS. The cells in four wells were infected with the diluted virus samples for 4 days in a humidified incubator at 37°C. The cells were observed for CPE under microscope.

#### 8.5. Antibody detection

The presence of IgG antibody specific for MERS-CoV or SARS-CoV-2 in the sera of infected and treated animals was determined using enzyme-linked immunosorbent assays (ELISA). The purified and inactivated MERS-CoV or SARS-CoV-2 antigen was diluted to final concentration of 100μg/ml in coating buffer (carbonate-bicarbonate buffer, pH 9.6). The diluted antigen (100μl) was coated to the wells of a Nunc-Immuno^™^ MicroWell^™^ 96 well solid plates (Sigma-Aldrich, MO, USA) and was incubated overnight at 4°C. After removing the coating buffer, the plate was washed twice by filling the wells with 400μl of washing buffer (0.05% tween 20 PBS (pH 7.4) containing 4% horse serum). To block the remaining protein-binding sites, 400μl of blocking buffer (PBS containing 4% skim milk) was added to the plate and incubated overnight at 4°C. The buffer was removed, and sera (100μl diluted in 1:64 in PBS) collected from treated mice on 14 days post treatment were added to the plate and incubated for 1hr at room temperature. The plate was washed 4 times with washing buffer. Goat anti-Mouse IgG Cross-Adsorbed Secondary Antibody HRP (Invitrogen, MA, USA) was diluted (1:5000) in blocking buffer, and 100 μL was added to each well and incubated for 1hr at room temperature. After washing the plate 4 times with the washing buffer, 100 μL of the TMB ELISA substrate (Mabtech, Nacka Strand, Sweden) was dispensed into the wells and incubated for 30min at 4°C. ABTS® Peroxidase Stop Solution (KPL, MD, USA) (100μl) was then added to the plate. The absorbance of each well was measured at 450nm using iMARK^™^ Microplate Absorbance Reader (Bio-Rad, CA, USA).

#### 8.6. Statistical analysis

Survival curves in both MERS-CoV and SARS-CoV-2 lethal infection models were evaluated for statistical significance using the Mantel-Cox test. Statistical analysis on tissue homogenate viral titres was performed using a two-way ANOVA with Dunnet’s multiple comparisons test to evaluate post-hoc differences between groups. Alpha was set at 0.05. Statistical analysis was performed using Prism8 (GraphPad Software, San Diego, CA, USA).

#### 8.7. Ethics approval

Protocols for the study of SARS-CoV-2 drug efficacy (202003-CNU-023) and MERS-CoV drug efficacy (CNU-01192) in mice was approved by the Internal Animal Use Committee at Chungnam National University (CNU). All the studies were approved and were conducted in accordance with the relevant legal guidelines and regulations prescribed by CNU, Republic of Korea.

## ACKNOLWEDGEMENTS

The authors wish to acknowledge the valuable contributions of Miguel Orlando Jara Gonzalez (University of Texas at Austin, College of Pharmacy) in the obtainment and review of background literature to support the premise of this work.

