## Supplementary Figure 1 for "Broad-Spectrum, Patient-Adaptable Inhaled Niclosamide-Lysozyme Particles are Efficacious Against Coronaviruses in Lethal Murine Infection Models"

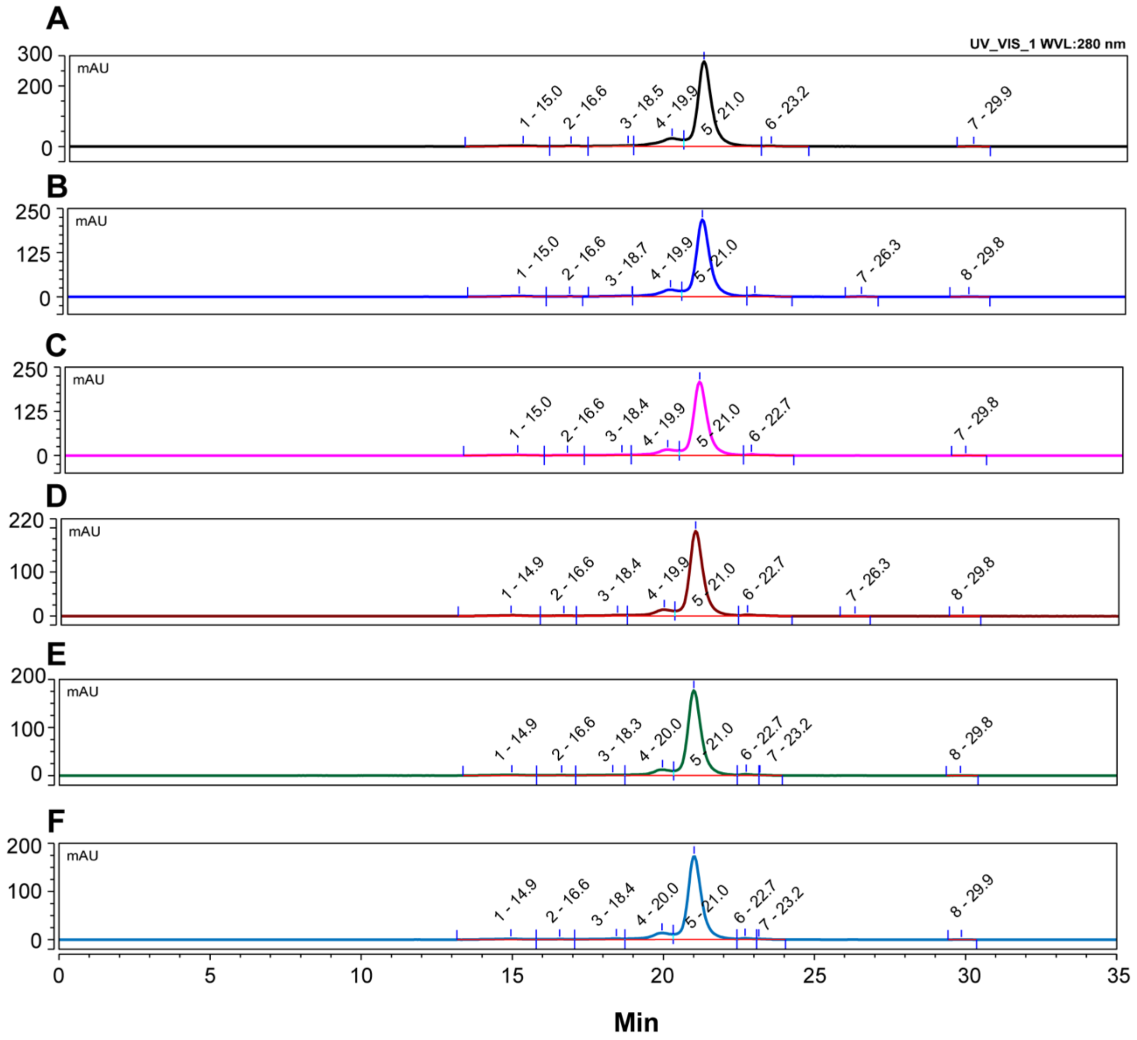


*Supplementary Figure 1: Spray drying (B) did not result in additional formation of soluble aggregates compared to the unprocessed control (A). NIC-hLYS reconstituted at a concentration of 25 mg/mL (C), 50 mg/mL (D), 75 mg/mL (E), and 100 mg/mL (F) was nebulized over the course of 2 minutes from an Aerogen Solo vibrating mesh nebulizer. This process did not result in additional aggregation compared to the unprocessed control.*
