## Supplementary Table 1 for "Broad-Spectrum, Patient-Adaptable Inhaled Niclosamide-Lysozyme Particles are Efficacious Against Coronaviruses in Lethal Murine Infection Models"

*Supplementary Table 1: Formulations developed for the production of inhalable NIC-hLYS particles and resulting particle size distributions*

| | % Components in 1% w/v feed suspension (prepared in 0.174 mg/mL histidine) | | | | | Reconstituted in ½ NS | | | Reconstituted in DI water | | | |
|---|---|---|---|---|---|---|---|---|---|---|---|---|
| Formulation | Micronized niclosamide (% w/w) | Human Lysozyme (% w/w) | Sucrose (% w/w) | Tween 80 (% w/w) | X90 diameter – dry powder (µm) | X50 diameter (µm) | X90 diameter (µm) | X99 diameter (µm) | X50 diameter (µm) | X90 diameter (µm) | X99 diameter (µm) | Decrease in monomer vs control (%) |
| 1 | 0.30 | 60.00 | 39.65 | 0.050 | 5.63 | 5.16 | 42.13 | 83.65 | 1.41 | 8.66 | 18.72 | 0.00 |
| 2 | 1.00 | 60.00 | 38.95 | 0.050 | 5.80 | 2.44 | 6.49 | 15.48 | 1.87 | 47.47 | 107.46 | 0.00 |
| 3 | 0.30 | 79.65 | 20.00 | 0.050 | 6.07 | 2.23 | 36.32 | 73.41 | 2.18 | 41.64 | 76.01 | 0.00 |
| 4 | 1.00 | 78.95 | 20.00 | 0.050 | 6.66 | 1.91 | 4.42 | 7.18 | 1.39 | 4.08 | 12.06 | 0.00 |
| 5 | 0.30 | 60.00 | 39.50 | 0.200 | 6.32 | 19.73 | 72.22 | 104.27 | 1.93 | 30.48 | 66.36 | 0.00 |
| 6 | 0.30 | 79.50 | 20.00 | 0.200 | 6.29 | 1.58 | 5.05 | 11.84 | 2.08 | 39.65 | 72.61 | 0.00 |
| 7 | 1.00 | 60.00 | 38.80 | 0.200 | 5.46 | 1.84 | 5.57 | 23.03 | 1.69 | 6.06 | 39.35 | 0.00 |
| 8 | 1.00 | 78.80 | 20.00 | 0.200 | 6.11 | 1.77 | 5.10 | 12.22 | 1.49 | 4.11 | 8.35 | 0.00 |
| 9 | 1.00 | 78.88 | 20.00 | 0.125 | 5.62 | 1.64 | 5.78 | 21.74 | 1.34 | 3.87 | 11.23 | 0.00 |
| 10 | 0.30 | 69.79 | 29.79 | 0.125 | 5.95 | 2.98 | 54.57 | 81.88 | 1.81 | 50.22 | 81.49 | 0.09 |
| 11 | 0.65 | 60.00 | 39.22 | 0.125 | 6.06 | 2.10 | 14.64 | 38.86 | 1.41 | 4.10 | 11.58 | 0.00 |
| 12 | 0.65 | 69.65 | 29.65 | 0.050 | 6.56 | 1.70 | 5.11 | 14.66 | 1.42 | 5.36 | 22.13 | 0.00 |
