## Supplementary Table 2 for "Broad-Spectrum, Patient-Adaptable Inhaled Niclosamide-Lysozyme Particles are Efficacious Against Coronaviruses in Lethal Murine Infection Models"

*Supplementary Table 2: Spray pattern analysis of varying concentrations of NIC-hLYS emitted from the VP7 Aptar® nasal spray device.*

| | | Spray Pattern 2 cm | | | Spray Pattern 5 cm | | |
|---|---|---|---|---|---|---|---|
| **Concentration (mg/mL)** | **Plume Angle (°)** | **Spray Area ($mm^2$)** | **Max Diameter (mm)** | **Min Diameter (mm)** | **Spray Area ($mm^2$)** | **Max Diameter (mm)** | **Min Diameter (mm)** |
| 10 | 42.3 ± 1.4 | 432 ± 8.7 | 24.4 ± 0.3 | 23.4 ± 0.2 | 1398 ± 123.7 | 49.6 ± 1.9 | 38.0 ± 2.4 |
| 25 | 45.5 ± 4.7 | 434 ± 9.9 | 26.7 ± 1.9 | 22.9 ± 0.2 | 1799 ± 151.5 | 58.8 ± 1.5 | 43.6 ± 1.8 |
| 50 | 58.6 ± 2.8 | 470 ± 3.8 | 28.5 ± 0.5 | 23.4 ± 0.25 | 2668 ± 1025.1 | 64.0 ± 14.0 | 55.1 ± 9.7 |
