## Supplementary Table 3 for "Broad-Spectrum, Patient-Adaptable Inhaled Niclosamide-Lysozyme Particles are Efficacious Against Coronaviruses in Lethal Murine Infection Models"

*Supplementary Table 3: Osmolality of varying concentrations of NIC-hLYS reconstituted in 0.45% sodium chloride*

| Concentration (mg/mL) | Osmolality (mOsmol/kg) |
|---|---|
| 10 | 156 ± 3 |
| 25 | 179 ± 4 |
| 50 | 208 ± 1 |
