## Supplementary Table 4 for "Broad-Spectrum, Patient-Adaptable Inhaled Niclosamide-Lysozyme Particles are Efficacious Against Coronaviruses in Lethal Murine Infection Models"

*Supplementary Table 4: Aggregation of hLYS before and after spray drying and nebulization*

| Sample | HMW (%) | Monomer (%) | Fragment (%) |
|---|---|---|---|
| hLYS unprocessed | 17.2 | 82.1 | 0.7 |
| NIC-hLYS-SD | 13.7 | 84.3 | 2.1 |
| 25 mg/mL nebulized | 14.1 | 84.2 | 1.7 |
| 50 mg/mL nebulized | 13.4 | 84.7 | 1.9 |
| 75 mg/mL nebulized | 12.5 | 85.8 | 1.6 |
| 100 mg/mL nebulized | 14.0 | 84.4 | 1.7 |
