## Supplementary Table 5 for "Broad-Spectrum, Patient-Adaptable Inhaled Niclosamide-Lysozyme Particles are Efficacious Against Coronaviruses in Lethal Murine Infection Models"

*Supplementary Table 5: Secondary structure of hLYS formulations as determined by FTIR*

| Sample | Anti-parallel β-sheet (%) (1618-1625 $cm^{-1}$) | Parallel β-sheet (%) (1633 – 1637 $cm^{-1}$) | α-helix (%) (1653 $cm^{-1}$) | Turns (%) (1678-1680 $cm^{-1}$) | Adj $R^2$ |
|---|---|---|---|---|---|
| hLYS unprocessed | 21.7 | 0.7 | 57.6 | 20.2 | 0.99 |
| hLYS-SD | 10.4 | 12.1 | 59.5 | 18.1 | 0.99 |
| NIC-hLYS-SD | 12.6 | 9.7 | 58.5 | 19.2 | 0.99 |
