## Supplementary Table 6 for "Broad-Spectrum, Patient-Adaptable Inhaled Niclosamide-Lysozyme Particles are Efficacious Against Coronaviruses in Lethal Murine Infection Models"

*Supplementary Table 6: Upper and lower constraints for mixture DoE*

| Component | Lower constraint (%w/w) | Upper constraint (%w/w) |
| --- | --- | --- |
| Micronized niclosamide | 0.3 | 1 |
| Human lysozyme | 60 | 80 |
| Sucrose | 20 | 40 |
| Polysorbate 80 | 0.05 | 0.2 |
| Histidine | Fixed at 0.174 mg/mL in feed solution | |
