## Supplementary Table 7 for "Broad-Spectrum, Patient-Adaptable Inhaled Niclosamide-Lysozyme Particles are Efficacious Against Coronaviruses in Lethal Murine Infection Models"

*Supplementary Table 7: D-optimal subset utilized for constratined mixtures DoE*

| Run | X1 (NIC-M) | X2 (hLYS) | X3 (sucrose) | X4 (polysorbate 80) | Dimension |
|---|---|---|---|---|---|
| 1 | 0.0030 | 0.6000 | 0.3965 | 0.0005 | 0 |
| 2 | 0.0100 | 0.6000 | 0.3895 | 0.0005 | 0 |
| 3 | 0.0030 | 0.7965 | 0.2000 | 0.0005 | 0 |
| 4 | 0.0100 | 0.7895 | 0.2000 | 0.0005 | 0 |
| 5 | 0.0030 | 0.6000 | 0.3950 | 0.0020 | 0 |
| 6 | 0.0030 | 0.7950 | 0.2000 | 0.0020 | 0 |
| 7 | 0.0100 | 0.6000 | 0.3880 | 0.0020 | 0 |
| 8 | 0.0100 | 0.7880 | 0.2000 | 0.0020 | 0 |
| 14 | 0.0100 | 0.7888 | 0.2000 | 0.0012 | 1 |
| 21 | 0.0030 | 0.6979 | 0.2979 | 0.0012 | 2 |
| 23 | 0.0065 | 0.6000 | 0.3922 | 0.0012 | 2 |
| 25 | 0.0065 | 0.6965 | 0.2965 | 0.0005 | 2 |
