## Supplementary Table 8 for "Broad-Spectrum, Patient-Adaptable Inhaled Niclosamide-Lysozyme Particles are Efficacious Against Coronaviruses in Lethal Murine Infection Models"

*Supplementary Table 8: Primer sequences used for the quantification of viral particles*

| Primer name | Sequence |
|---|---|
| MERS N3 Forward | GGG TGT ACC TCT TAA TGC CAA TTC |
| MERS N3 Reverse | TCT GTC CTG TCT CCG CCA AT |
| MERS N3 probe | 5'FAM-ACC CCT GCG CAA AAT CGT-BHQ1 3' |
| SARS-CoV-2 Forward | CAC ATT GGC ACC CGC AAT C |
| SARS-CoV-2 Reverse | GAG GAA CGA GAA GAG GCT TG |
| SARS-CoV-2 Probe | 5'FAM-ACT TCC TCA AGG AAC AAC ATT GCC A-BHQ1 3' |
